# A Comparison Study between Autism Spectrum Disorder and Typically Control in Graph Frequency Bands Using Graph and Triadic Interaction Metrics

**DOI:** 10.1101/2021.11.19.469268

**Authors:** Alireza Talesh Jafadideh, Babak Mohammadzadeh Asl

## Abstract

Many researchers using many different approaches have attempted to find features discriminating between autism spectrum disorder (ASD) and typically control (TC) subjects. In this study, this attempt has been continued by analyzing global metrics of functional graphs and metrics of functional triadic interactions of the brain in the low, middle, and high-frequency bands (LFB, MFB, and HFB) of the structural graph. The graph signal processing (GSP) provided the combinatorial usage of the functional graph of resting-state fMRI and structural graph of DTI. In comparison to TCs, ASDs had significantly higher clustering coefficients in the MFB, higher efficiencies and strengths in the MFB and HFB, and lower small-world propensity in the HFB. These results show over-connectivity, more global integration, and probably decreased local specialization in ASDs compared to TCs. Triadic analysis showed that the numbers of unbalanced triads were significantly lower for ASDs in the MFB. This finding may show the reason for restricted and repetitive behavior in ASDs. Also, in the MFB and HFB, the numbers of balanced triads and the energies of triadic interactions were significantly higher and lower for ASDs, respectively. These findings may reflect the disruption of the optimum balance between functional integration and specialization. All of these results demonstrated that the significant differences between ASDs and TCs existed in the MFB and HFB of the structural graph when analyzing the global metrics of the functional graph and triadic interaction metrics. In conclusion, the results demonstrate the promising perspective of GSP for attaining discriminative features and new knowledge, especially in the case of ASD.

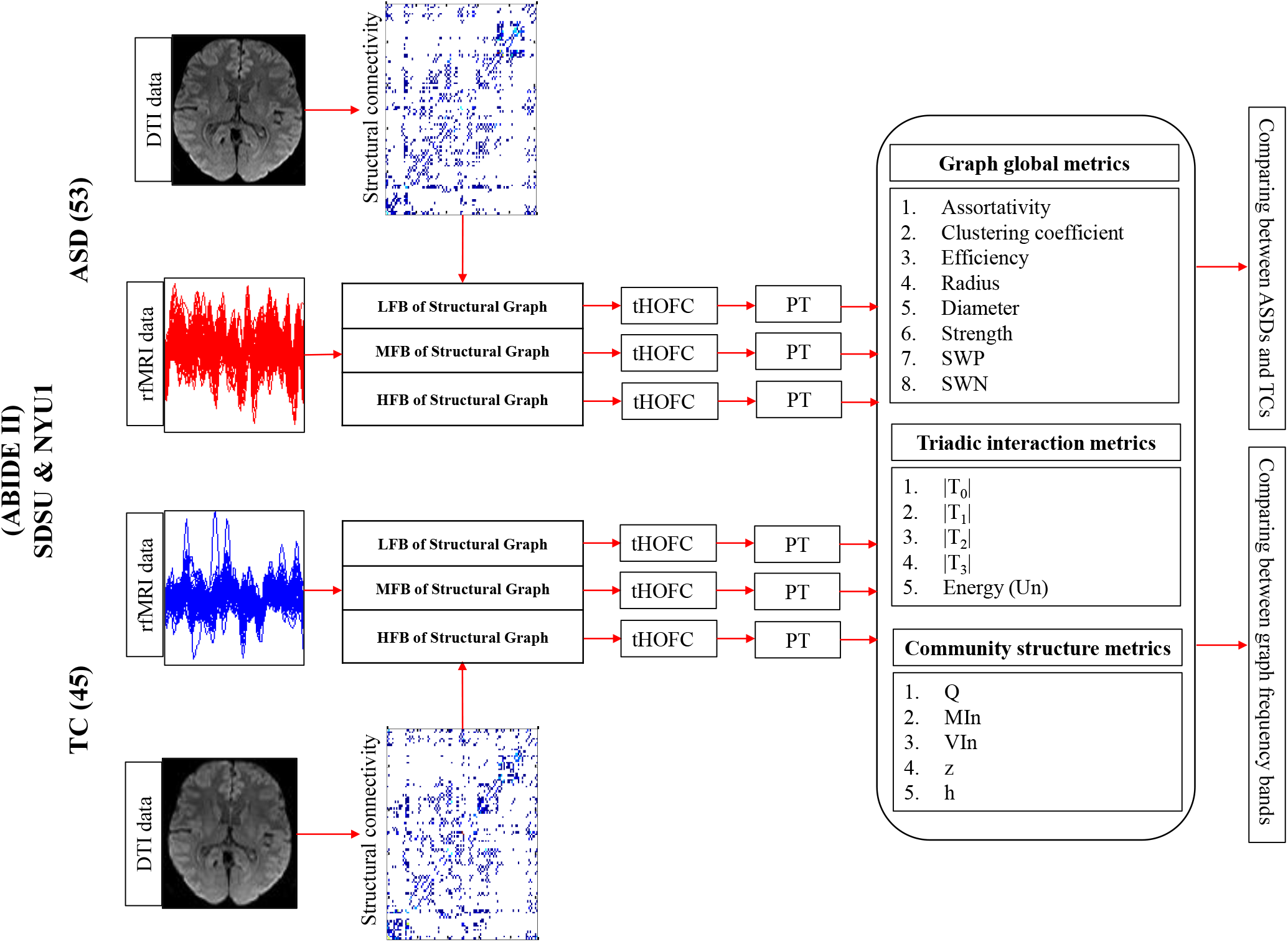
Graphical Abstract.

## 1. Introduction

Autism spectrum disorder (ASD) is a neurological and neurodevelopmental disorder characterized by impaired social communication, restricted and repetitive interests, behavior, and activities (American Psychiatric Association, 2013). Because of the rapid prevalence and heterogeneous etiology of ASD, a variety of researches have been devoted to finding biomarkers and features discriminating ASDs from typically controls (TCs) (Goldani et al., 2014; Woo and Wager, 2015; Drysdale et al., 2016; Li et al., 2017).

One of the popular approaches for finding reliable and quantifiable biomarkers is studying the functional behavior of the brain through resting-state functional magneto resonance imaging (rfMRI) data (Grecucci et al., 2017; Uddin et al., 2013; Ha et al., 2015; Mash et al., 2018; Mash et al., 2019; Rashid et al., 2018). However, the inherent complexity of the human brain makes its functional behavior study a challenging issue (Bullmore and Sporns, 2009). To handle this complexity, the brain is usually modeled by a graph whose vertices are regions of interest (ROI)s of the brain and edges represent the functional connectivity (FC) between regions of interest (ROIs) (De Vico Fallani et al., 2014; Song et al., 2019). The most commonly used metrics of the graph can be classified into two groups: local and global metrics (Fornito et al., 2016; Farahani et al., 2019). The earlier ones describe the network behavior at the ROI level and offer discriminative features at this level. The latter ones describe the properties governing the entire brain network.

The ASD leads to abnormal FCs (Mash et al., 2019; Rashid et al., 2018), which in turn change the graph metrics. Thus, these changes can be revealed and regarded as candidate biomarkers (Hull et al., 2016). Di Martino and colleagues (Di Martino et al., 2013) found that cortical and subcortical ROIs of ASDs had more connections (degree) than TCs. Redcay E and colleagues (Reday E et al., 2013) reported the greater betweenness centrality, a metric indicating the ROI centrality, for ASDs compared to TCs when studying the right parietal region of the default mode network (DMN). Both of these findings represent the overconnectivity in ASDs compared to TCs (Hull et al., 2016). Rudie and colleagues (Rudie et al., 2013) reported decreased local and increased global efficiency in adolescents with ASD. Itahashi and colleagues (Itahashi et al., 2014) found reduced characteristic path length and clustering coefficient when studying adult ASDs in comparison to TCs. The results of these two papers show the increased randomness of functional networks for ASDs’ brains (Rudie et al., 2013; Keown et al., 2017). Keown and colleagues (Keown et al., 2017) reported findings at the global level. Their analysis showed globally reduced cohesion but increased dispersion of networks. Cohesion quantified how well nodes from a normative network grouped within the community structure. Dispersion quantified how distributed nodes from a network were in the community structure. These results demonstrated impaired network integration and segregation of ASD subjects (Keown et al., 2017). Zhou and colleagues (Zhou et al., 2014) performed a small-world network analysis using the FC and did not find any differences between ASD and TC children. Altogether, impaired functional organizations of the autistic brain were found in both local and global metrics of the graph (Farahani et al., 2019).

In most of the graph metrics and methods of FC analysis, the dyadic interactions are exploited. However, the human brain is one of the most complex networks in the world and the interaction of two ROIs is not independent of the rest of the ROI-ROI interactions. Thus, a higher level of interactions exists in the brain network and investigation of them can provide new valuable findings. Recently, analyzing such higher-level interaction was carried out by Moradimanesh and colleagues (Moradimanesh et al., 2021). The authors analyzed the brain triadic interactions to compare ASD and TC groups. For analyzing the triadic interactions, three ROIs and their interactions with each other were considered. The ROIs interactions was FC measured by Pearson correlation. The authors investigated four types of triad interactions, including strongly balanced T3: (+ + +), weakly balanced T1: (+ – –), strongly unbalanced T2: (+ + –), and weakly unbalanced T0: (– – –). The “+” and are the sign of Pearson FC between ROIs. They found that balanced and unbalanced triads were over-presented and under-presented in both ASD and TC groups, respectively. Also, they observed that the energy of the salience network (SN) and the default mode network (DMN) were lower in ASD, probably indicating the difficulty of flexible behavior.

The brain signals not only change over time but also change over the brain topology. The topology domain is irregular. As a result, the signal changes in the topological domain cannot be captured by the Fourier transform of the time domain. To address this problem, the field of graph signal processing (GSP) has been recently emerged attempting to develop methods for capturing frequencies of the topological domain (Shuman et al., 2013; Ortega et al., 2018). The GSP needs two elements: graph and signal. The graph represents the underlying structure of the signal and the signal is brain activity mounted on the graph. Using these elements and graph Fourier transform (GFT), one can deal with topological frequencies. The GFT is one of the GSP tools for working with topological frequencies. Recently, the GFT has been used in two studies of autism to classify ASDs and TCs (Brahim and Farrugia, 2020; Itani and Thanou, 2021). Brahim and Farrugia (Brahim and Farrugia, 2020) used the structural connectivity matrix of the Human Connectome Project (HCP) to obtain the graph of GSP. This matrix had been attained by averaging over 56 TC subjects. The authors used this averaged structure as the underlying structure of all studied ASD and TC subjects. Some statistical metrics including standard deviation (SD), mean, variance, and Kurtosis were computed for each ROI. These computed metrics were considered as the signal of GSP. These signals and their GFT were used as features. The authors found that the feature with the best classification performance was the GFT of SD. Itani and Thanou (Itani and Thanou, 2021) used the inverse of the distance between ROIs as weights to construct the graph of GSP. The authors proposed a framework using both GFT and spatial filtering method (SFM) to construct a subspace extracting discriminative features between ASDs and TCs. The classification using the decision tree demonstrated the superiority of their proposed method performance in comparison to other state-of-the-art ones.

In this study, the graph of GSP was attained by SC matrix and the signal was rfMRI data. The SC matrix was computed using the diffusion tensor imaging (DTI) data. Using GFT, graph frequency modes and filters were obtained. Then, rfMRI data was filtered to have data in three independent graph frequency bands, i.e., low, middle, and high-frequency bands. Hereinafter, these bands are abbreviated as LFB, MFB, and HFB, respectively. For each band, the FC matrix was computed using the filtered data. From these FC matrices, some of the graph global metrics and triadic interaction metrics were computed and compared between ASD and TC groups Also, for each of the ASD and TC groups, the behavior of metrics were compared between three aforementioned frequency bands. By doing so, we hope to reveal significant differences between ASDs and TCs.

The rest of the paper is organized as follows: Section 2 describes the dataset, preprocessing methods, and brain ROIs and networks. This section is continued by explaining the GFT and graph filtering and then describing the scenarios. The last subsection of this section is devoted to statistical evaluations. Section 3 provides the results. The discussion about the results and conclusions are respectively given in sections 4 and 5.

## 2. Materials and methods

### 2.1. Participants and data acquisition

In this study, the SDSU and NYU1 datasets of ABIDE II database were exploited. In the NYU1 dataset, only 57 subjects (33 ASDs, 24 TCs) had both DTI and fMRI data. Thus, only these subjects were selected for the next step. Both datasets had mostly male subjects for both ASD and TC groups. Accordingly, only the male subjects were analyzed.

For both datasets, two groups were matched on age, handedness, and head motion while showing significant differences in social responsiveness scale (SRS). The sample characteristics are listed in Table 1. This information is for subjects whose data were survived in preprocessing step. The SRS values of one SDSU subject (ASD) and three NYU1 subjects (ASD=2, Tc=1) were missing. The framewise displacement (FD) is explained in subsection 2.2.1.

**Table 1.**
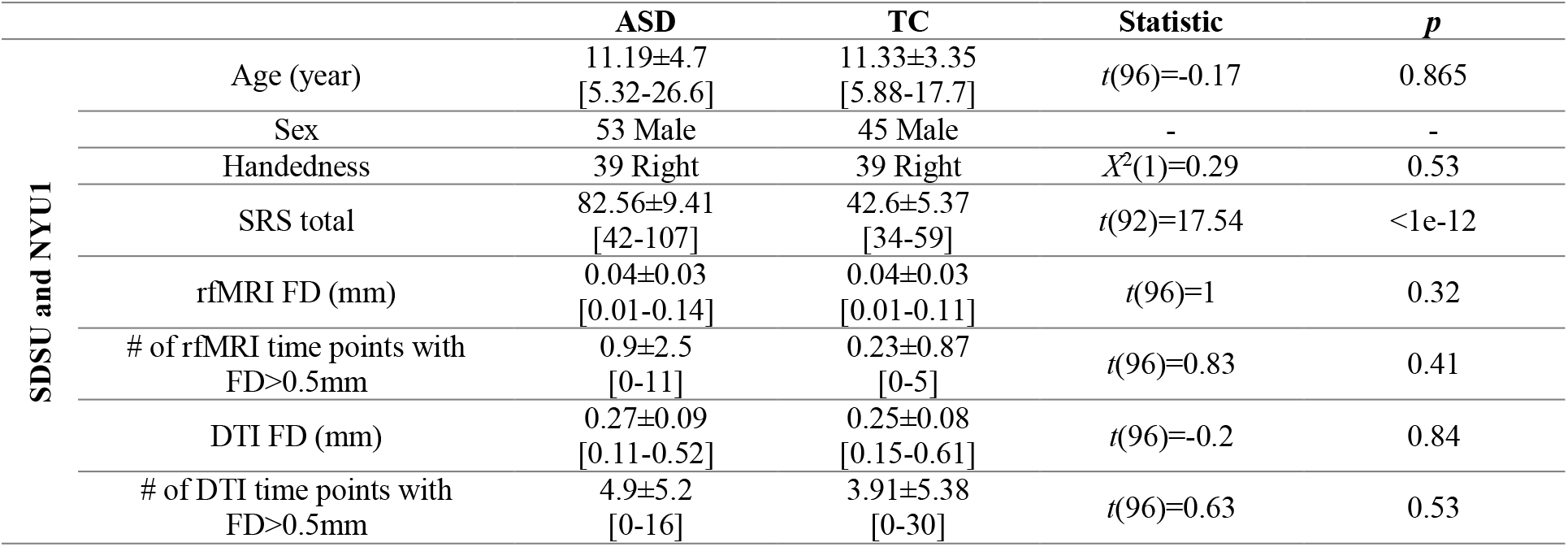
Sample characteristics.

|  | ASD | TC | Statistic | *p* |
| --- | --- | --- | --- | --- |
| Age (year) | 11.19±4.7<br>[5.32-26.6] | 11.33±3.35<br>[5.88-17.7] | *t*(96)=-0.17 | 0.865 |
| Sex | 53 Male | 45 Male | - | - |
| Handedness | 39 Right | 39 Right | $\chi^2(1)=0.29$ | 0.53 |
| SRS total | 82.56±9.41<br>[42-107] | 42.6±5.37<br>[34-59] | *t*(92)=17.54 | <1e-12 |
| rfMRI FD (mm) | 0.04±0.03<br>[0.01-0.14] | 0.04±0.03<br>[0.01-0.11] | *t*(96)=1 | 0.32 |
| # of rfMRI time points with<br>FD>0.5mm | 0.9±2.5<br>[0-11] | 0.23±0.87<br>[0-5] | *t*(96)=0.83 | 0.41 |
| DTI FD (mm) | 0.27±0.09<br>[0.11-0.52] | 0.25±0.08<br>[0.15-0.61] | *t*(96)=-0.2 | 0.84 |
| # of DTI time points with<br>FD>0.5mm | 4.9±5.2<br>[0-16] | 3.91±5.38<br>[0-30] | *t*(96)=0.63 | 0.53 |

The data were collected using (GE 3T MR750 scanner, 3T Siemens Allegra) with (eight-channel, eight-channel) head coils. High-resolution structural images were acquired according to (SPGR, TFL) standards’ T1-weighted sequences and with TR/TEs of (8.136/3.172 ms, 3.25/3.25 ms), flip angles of (8°, 7°), fields of view (FOVs) of (256×256 mm^2^, 256×256 mm^2^), resolutions of (1 mm^3^, 1.73 mm^3^), and numbers of slices of (172, 128). The rfMRI datasets were collected using standard (gradient echo-planar imaging (EPI), EPI) sequences with TR/TEs of (2000/30 ms, 2000/15 ms), flip angles of (90°, 90°), FOVs of (220×220 mm^2^, 240×240 mm^2^), matrix sizes of (64×64, 80×80), pixel spacing sizes of (3.4375×3.4375 mm^2^, 3×3 mm^2^), slice thicknesses of (3.4 mm, 4 mm), slice gaps of (0 mm, 0 mm), axial slices of (42, 33), and total volumes of (180, 180). The DTI protocols were based on EPI sequences. The DTI images consisted of (61, 64) weighted diffusion scans with *b* values of (1000 sec/mm^2^, 1000 sec/mm^2^) and one unweighted diffusion scan with *b* values of (0 sec/mm^2^, 0 sec/mm^2^). The other parameters of data recording included TR/TEs of (8500/78 ms, 5200/78 ms), FOVs of (128×128 mm^2^, 192×192 mm^2^), slice in-place resolutions of (1.875×1.875 mm^2^, 3×3 mm^2^), slice thicknesses of (2 mm, 3 mm), and the numbers of axial slices of (68, 50).

### 2.2. Data preprocessing

#### 2.2.1. rfMRI data preprocessing and time series extraction

The SPM^1^ and AFNI^2^ software packages and personal codes were used for preprocessing the fMRI data. To prevent T1- equilibration effects, the first five volumes were ignored. The data were corrected for slice-timing by considering the middle slice as a reference, motion artifact. The data were despiked through the AFNI’s 3dDespike function. Then, the spm_coreg function was applied to coregister the T1 image and functional images. The images warped to the standard Montreal Neurological Institute (MNI) space using a template created by nonlinear registration of 152 T1-weighted images. A Gaussian kernel with a 5-mm full width at half maximum (FWHM = 5 mm) was employed for spatial smoothing.

For more noise reduction of data, six motion parameters and their first temporal derivatives and one principal component of cerebrospinal fluid (CSF) and white matter (WM) signals were considered 14 confounds and regressed out from all voxels’ time series. These principle components of CSF and WM explained more than 99% variance of time series and were considered as confounds (Caballero-Gaudes and Reynolds, 2017). Then, removing the linear, quadratic, and cubic trends from the time series and applying a band-pass fifth-order Butterworth filter (0.01-0.1 Hz) on the time series were carried out.

To detect the volumes corrupted by motion artifacts, the FD was computed. The FD was defined as the square root of the sum of squares of differences existing in motion parameters of two consecutive time points. The head was considered a sphere with a radius of 50mm to change the values of rotational parameters from radian to millimeter (Power et al., 2012). The time points with FD >0.5 mm and the two subsequent time points were removed (Mash et al., 2019). After censoring, as with Mash and colleagues (Mash et al., 2019), the fragments of time series with less than 10 consecutive time points were removed. After censoring, the data was used if more than %75 of its time points were preserved from censoring. The TC subject with ID = 28852 of the SDSU dataset was rejected. The number of subjects of ASD and TC without censoring time points were 42 and 36, respectively, (χ2(1) = 0.0009, *p* = 0.97). There was no significant difference between ASD and TC in terms of the number of censored time points (t(94) = 0.85, *p* = 0.39). Given that the maximum number of censored time points was 20 and the first five volumes were removed for preventing from T1-equilibrium effect, the 155 time points were used for further analysis.

#### 2.2.2. DTI preprocessing and structural connectivity matrix

Preprocessing of DTI data and computation of structural connectivity matrix were carried out using the ExploreDTI software (Leemans et al., 2009). Firstly, data preparation and quality assessment were performed in ExploreDTI. After that, diffusion data were corrected for subjects’ motion and Eddy currents. Then, deterministic fiber tracking was performed for tractography. The results of tractography and ROIs of the studied atlas were employed for the calculation of the structural connectivity matrix. The structural connectivity between two ROIs were defined as the number of tracts existing between them. In the next step, each row of the matrix was divided by the sum of its elements. By doing so, the value of entry (*i,j*) expressed the connectivity probability from region *i* to region *j*, which was not equal to that from region *j* to region *i*. Thus, the connectivity matrix, which is also called the adjacency matrix **A**, was not symmetric. As the final step, the symmetric adjacency matrix **A’** was obtained as **A’** = (**A** + **A^T^**)/2 where.^T^ denotes the transpose operator. The FDs of DTI data are summarized in Table 1. In ExploreDTI, the REKINDLE (Robust Extraction of Kurtosis INDices with Linear Estimation) (Tax C.M.W. et al, 2015) detects and removes the outliers.

#### 2.3. Brain ROIs and networks

In this study, the Schaefer atlas (Schaefer et al., 2018) with 100 ROIs was employed. This atlas is purely based on functional time series and respects the boundaries of 7 Yeo-Krienen functional atlas (Yeo et al., 2011). For NYU1, one ASD subject was rejected. More than 10% of this subject’s ROIs had no structural connections with the rest of the ROIs. By averaging the time series of ROI voxels, one signal was obtained for each ROI.

#### 2.4. Graph frequency bands (GFBs)

The frequency content of the graph signal is defined according to the signal changes across connected vertices at a given time point. In low frequency, connected vertices show similar signals (representing alignment). In high frequency, the variability of the connected vertices signals is high compared to each other (representing liberality). In liberality, the vertices (brain ROIs) show less respect to their underlying connectivity structure. By approaching from low frequency to high frequency, the graph signal behavior changes from alignment to liberality (Fig 1).

**Fig 1.**
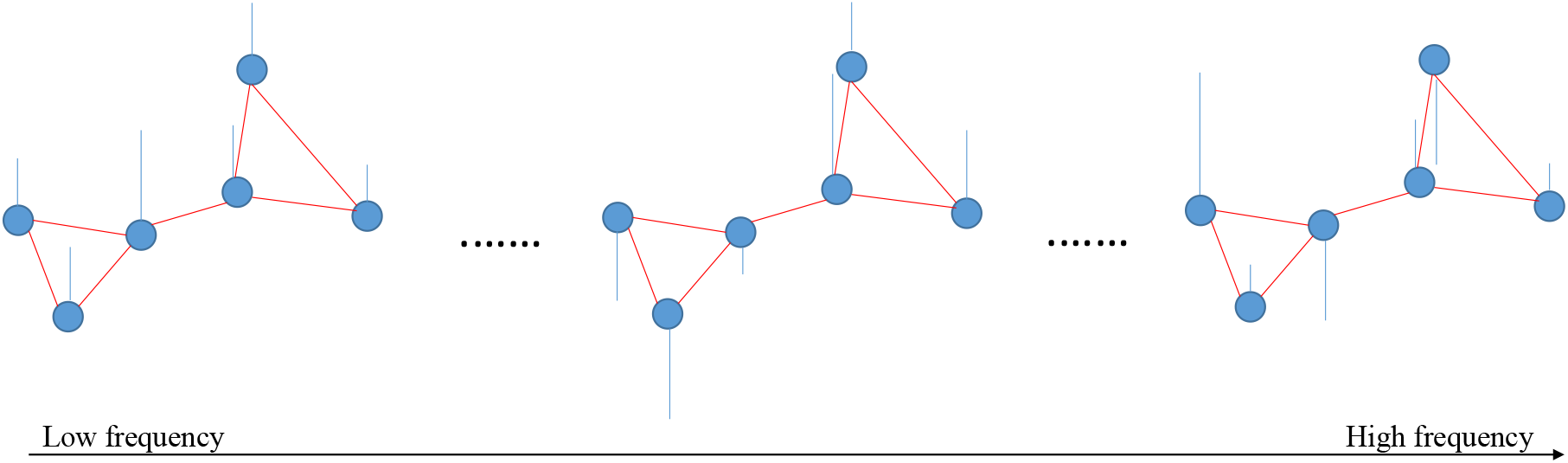
Simple representation of low and high graph frequency signals at a given time point. Blue circles, red and blue lines are vertices, edges, and signals, respectively.

The graph frequencies are defined using the combinatorial Laplacian matrix **L** ϵ ℝ^*N×N*^, (Shuman et al., 2013), as follows:

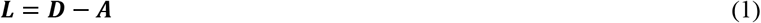

where **D** is a diagonal matrix and its *k*^th^ diagonal element represents the degree of *k*^th^ vertex i.e., 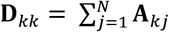. The eigendecomposition of **L** provides the **V** and **Λ**, which are the eigenvector matrix and diagonal eigenvalues matrix, respectively. The eigenvectors represent graph frequency modes and are used for GFT. The GFT of brain signal **x** ϵ ℝ^*N*×1^ is obtained as

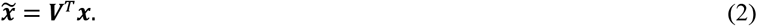

The inverse GFT (IGFT) of 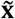 is attained by

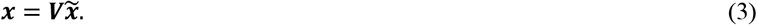

It is noteworthy that the eigenvector corresponding to the larger eigenvalue shows more variance and can pass higher graph frequencies (Huang et al., 2018). This means that higher frequency modes can transform brain signals of higher variance to the graph frequency domain and inversely transform the higher frequency information from the frequency domain to the brain topological domain.

The graph signal can be filtered at the frequency domain and, then, got IGFT to have graph filtered signal. The graph filtering process can be formulated as

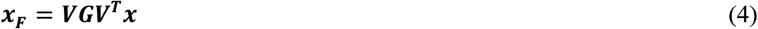

where **G** is a diagonal filtering matrix. This study considered 1 for the diagonal elements corresponding to the desired frequency modes and 0 for the rest of the modes.

In this study, the first 33 and the last 33 frequency modes formed the LFB and HFB and the rest 34 modes formed the MFB. Using the “(4)”, the fMRI data was filtered to have data in graph LFB, MFB, and HFB. Then, for each subject, the FC matrix was computed in each frequency band.

#### 2.5. Functional connectivity matrix

Recently, high-order FC (HOFC) methods have been used for obtaining the FC matrices (Zhou et al., 2020; Zhang et al., 2017). In this study, the topographical profile similarity-based HOFC (tHOFC) was used (Zhou et al., 2020). In this method, the Pearson correlations of a given ROI with the rest of the ROIs are computed to be obtained the low-order FC (LOFC) profile of the given ROI. This is performed for each of the ROIs. Then, tHOFC is measured as the similarity of LOFC profiles between each pair of brain regions. The tHOFC can offer supplementary information to the conventional LOFC and introduce more differences between groups (Zhou et al., 2020; Zhang et al., 2019).

To remove spurious connections and to obtain sparsely connected matrices, proportional thresholding was applied on FC matrices (Van den heuvel et al., 2017). This type of thresholding provides an equal number of connections and connection density for all subjects of both ASD and TC groups (Van den heuvel et al., 2017; Van wijk et al., 2010). Applying this thresholding preserves PT% of the strongest connections. In this study, the PT was changed from 10 to 50 with a step of 5. The PTs < 40 resulted in the fragmented graph for some subjects in the MFB and HFB. The PTs of 40 and 45 offered similar results. To have more sparse matrices, the PT=40 was selected as the final threshold value. The sparse connected matrices were used for scenarios of this study.

#### 2.6. Scenarios

All of the global metrics of the graph were calculated by GraphVar software (Kruschwitz et al., 2015). The analysis of triadic interactions was performed by personal codes. In each GFB, the metrics were statistically compared between ASD and TC groups. Also, for a given group, the metrics were statistically compared between LFB, MFB, and HFB of the graph.

##### 2.6.1. Global metrics of graph

The studied graph global metrics were assortativity, clustering coefficient, efficiency, radius, diameter, strength, SWN, and small-world propensity (SWP) (Fornito et al., 2016).The **assortativity** coefficient is a Pearson correlation between the degrees of all vertices on two opposite ends of a link (degree of a vertex is the number of edges connected to it). This coefficient takes values between −1 and 1. The positive values show that vertices with similar degrees tend to connect. The negative assortativity value of a graph states that the tendency of vertices with larger degrees is to connect to vertices with smaller degrees.

The **Clustering coefficient** is the fraction of triangles around a vertex and takes values between 0 and 1. The value 1 means that connected vertices to a given vertex are also connected. Fewer connections in the neighborhood of a given vertex lead to less value of the clustering coefficient. In this study, the clustering coefficient value averaged over ROIs was reported for each subject.

The **efficiency** is the average inverse shortest path length in the graph. In a weighted graph, the shortest path length between two vertices is equal to the minimum sum of edge weights existing between them. This metric measures the efficiency of information exchanging in the graph.

Among all the maximum distances between a vertex to all other vertices, the **radius**/**diameter** is the minimum/maximum one. The **strength** is the sum of weights of adjacent edges to the vertex. In this study, the strength value averaged over ROIs was reported for each subject.

The **SWN**s are graphs between lattice and random graphs, showing both high clustering coefficient and low average shortest path length (Lavg). Such high and low properties make SWNs suitable for representing real-world networks such as brain networks. In the brain, the high clustering coefficient indicates the functional specialization (processing information within each cluster of brain regions), and low L_avg_ represents the efficient functional integration of brain sub-graphs (Bassett and Bullmore, 2017; Fornito et al., 2016). In a formal analysis, it should be determined which clustering coefficient is “high” and which path length is “low” in a network. To address this problem, the normalized clustering coefficient and L_avg_ are used for the calculation of SWN metric. Thus, the SWN metric (σ) is computed as

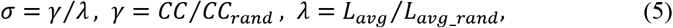

where *CC_rand_* and *L_avg_rand_* are computed in an ensemble of randomized surrogate networks. In a small-world network, a comparable path length and a higher clustering coefficient than a random network are expected (*λ*~1, *γ* > 1, *σ* > 1) (Fornito et al., 2016).

The definition of “(5)” has some limitations. Firstly, a network may have *σ* > 1 even when its L_avg_ is much greater than L_avg_ of a random graph. This means that a small world network always has *σ* > 1, but a network with *σ* > 1 is not necessarily a small world. Secondly, the *σ* is sensitive to the density of the network. The denser networks have smaller *σ* even if they are generated from an identical small-world model (Bassett and Bullmore, 2017). To overcome these and other limitations, Muldoon and colleagues developed SWP by employing the *CC* and of both lattice and random networks (Muldoon et al., 2016). The SWP measure *ϕ* is computed as

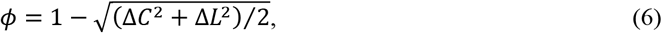

where

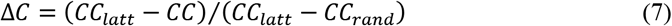

and

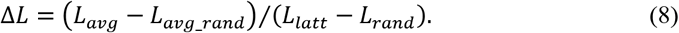

The authors set the Δ*C* and Δ*L* to 1/0 when they were larger/smaller than 1/0. By doing so, it is guaranteed that the *ϕ* is bounded in the range [0,1]. Networks with 0.4 < *ϕ* ≤ 1 are considered small-world (Bassett and Bullmore, 2017).

##### 2.6.2. Triadic interactions and their metrics

In this study, as with Moradimanesh and colleagues (Moradimanesh et al., 2021), four types of triads were analyzed in the LFB, MFB, and HFB of the structural graph. These triads were strongly balanced T3: (+ + +), weakly balanced T1: (+ – –), strongly unbalanced T_2_: (+ + –), and weakly unbalanced T0: (– – –). Two metrics were used for comparing triadic interactions between ASDs and TCs, including the number of the triad which is also called the frequency of triad (|T_i_|), where i=0,1,2,3, and the energy of the whole-brain network (Un). The Un is defined as

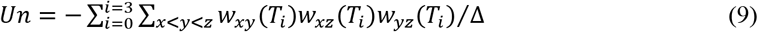

where x, y, and z indicate the ROIs of triad T_i_, *w_xy_* is the FC value, and Δ is the total number of triads of the brain.

In the brain network, it was shown that balanced/unbalanced triads were over-/under-presented (Moradimanesh et al., 2021). This means that the frequencies of balanced/unbalanced triads of the real network were more/less than those of random networks. The surprise value (S(T_i_)) is a metric providing positive/negative values for over-/under-presented of triads (Moradimanesh et al., 2021). This metric is defined as

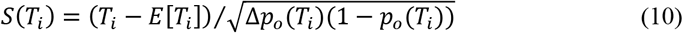

where *E*[*T_i_*] = *p_o_*(*T_i_*)Δ is the expected number of triads T_i_ and Δ is the total number of triads in the random network. The *P_o_* (*T_i_*) is the ratio of the number of triad Ti to the total number of triads in the random network. The random network had the same number of positive and negative links as the real brain network and the signs of the links were randomly assigned to the existing links. In this study, the S(T_i_) was only used to provide information for the over-/under-presented behavior of triads in the LFB, MFB, and HFB of the graph.

#### 2.7. Statistical analysis

The general linear model (GLM) was applied with age, diagnosis, and site variables as between covariate, between factor, and nuisance covariate, respectively. The diagnosis variable was set to 1 and 0 for ASD and TC subjects, respectively. The site variable was set to 1 and 0 for SDSU and NYU1 data, respectively. The nuisance covariate means that the site effect was regressed out from the dependent variable (studied metrics) before performing GLM. The GLM was for between-group comparisons. For a given group, the comparison between GFBs was performed using the t-test.

The non-parametric permutation testing with 400 repetitions was performed to correct the results. For between groups/GFBs comparisons, in each repetition, the measure value of each subject was randomly assigned to one group/GFB so that the final numbers of subjects of groups/GFBs were equal to the numbers of original groups/GFBs. Then, the difference of measure between two groups/GFBs was calculated. This process was repeated 400 times to obtain a distribution of measure difference. The probability of original between-group/between-GFB difference was calculated as its percentile position in this distribution.

## 3. Results

In this section, the degree of freedom of all *t* values is 96 and the *p-values* are corrected.

### 3.1. Results of graph metrics

The results of graph global metrics are shown in Fig 1. For **assortativity**, there was seen no significant difference between groups and between GFBs. This meant that both groups had similar resistance against failures in the main components of their network (i.e., their ROIs and edges) (Farahani et al., 2019). Such similarity was not sensitive to graph frequencies.

For **clustering coefficient**, the ASDs took higher values than TCs in all GFBs, particularly in the MFB and HFB. However, the dominancy of ASDs was significant only in the MFB (*t = 2.11, p = 0.04*). Both groups showed a similar pattern of clustering coefficient changes across GFBs i.e., decreased clustering coefficient by moving from LFB to MFB and increased clustering coefficient by moving from MFB to HFB. For TC group, the clustering coefficient values in the MFB and HFB were significantly lower than clustering coefficient value in the LFB (*t = −3.11, p = 0.005* for MFB; *t = −1.93, p = 0.002* for HFB)

For **efficiency**, the ASD group had similar behavior across the GFBs. In contrast, the efficiency of TC group was decreased by increasing the graph frequencies. As a result, significant differences between ASDs and TCs were seen in the MFB (*t = 2.11, p = 0.04*) and HFB (*t = 1.93, p = 0.04*). Also, there were significant differences between MFB and LFB (*t = −1.73, p = 0.03*) and between HFB and LFB (*t = −2.02, p = 0.02*) for TC group.

For the **radius** metric, there were no significant results. The common pattern for the two groups was increasing the radius by increasing the graph frequencies.

For **diameter** metric, similar behavior to radius metric was seen. However, the differences between GFBs reached a significant level. The diameter of HFB was significantly larger than that of LFB for both groups (*t = 3.74, p = 0.0005* for ASDs; *t = 3.22, p = 0.002* for TCs). For ASDs, another significant result was found between HFB and MFB (*t = 2.23, p = 0.03*).

For **strength**, ASDs had significantly larger values than TCs in the MFB (*t = 1.94, p = 0.047*) and HFB (*t = 1.93, p =0.049*).

**Fig. 1.**
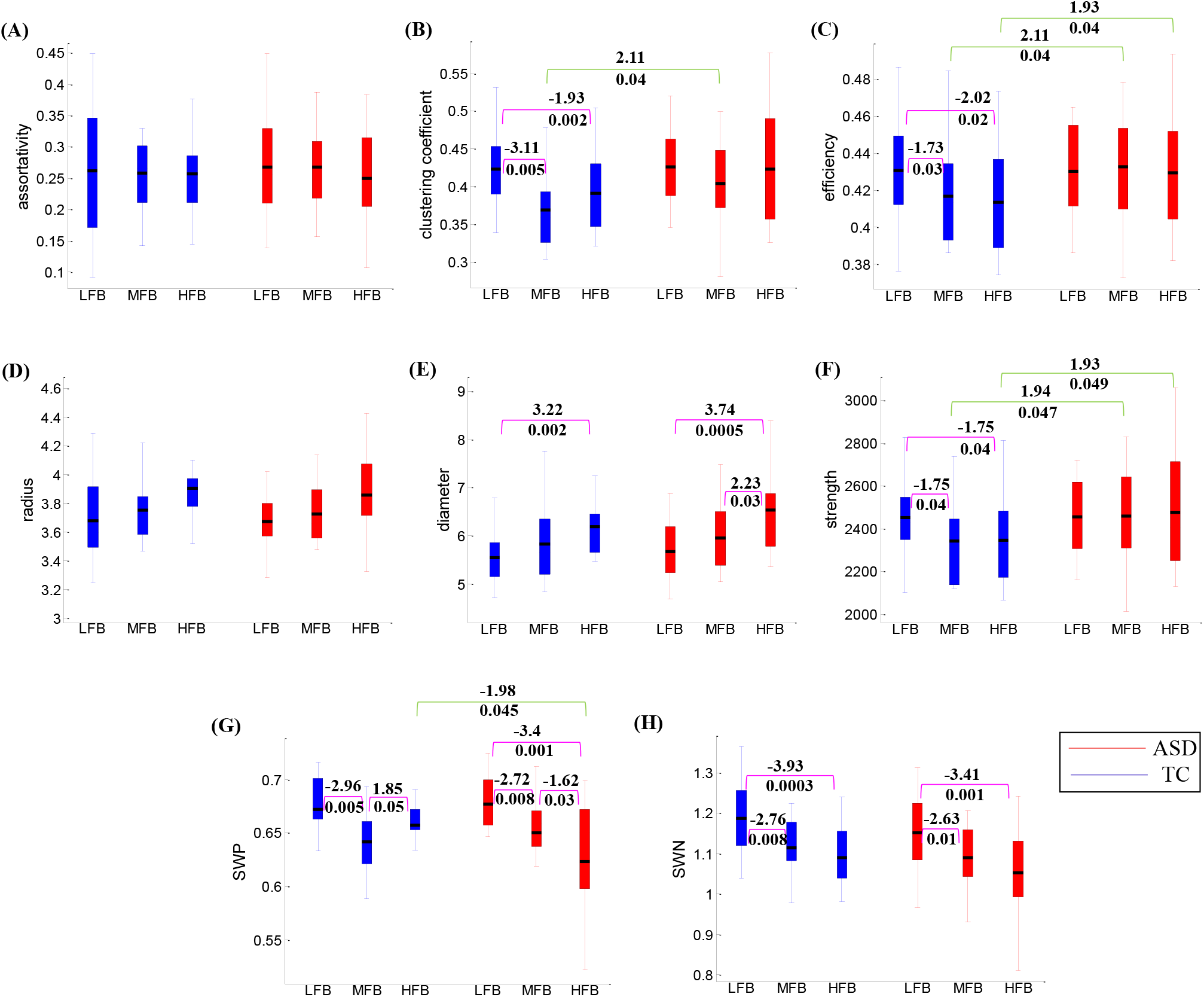
Comparing the graph global metrics between groups and between graph frequency bands. The results are for **(A)** Assortativity, **(B)** Clustering coefficient, **(C)** Efficiency, **(D)** Radius, **(E)** Diameter, **(F)** Strength, **(G)** SWP, and **(H)** SWN metrics. The green and pink color lines are used to show the results of between groups and between frequency bands, respectively. The values above and under the lines are statistical t and corresponding corrected p values, respectively. The aforementioned values are only provided for significant results (*p – 0.05*).

Strength values of the ASD group didn’t show changes across GFBs. The strength values of the TC group were similar to each other in the MFB and HFB and were significantly lower than the strength value in the LFB (*t = −1.75, p = 0.04* for both MFB and HFB).

Both **SWP** and **SWN** are metrics for investigation of small-world properties of complex networks. However, the SWP is the updated and modified one (Fornito et al., 2016; Bassett and Bullmore, 2017). The between groups pattern, which was common in both SWP and SWN, was the lower small-world property of ASDs than TCs in the HFB. This pattern reached to the significant level when using SWP metric (*t = −1.98, p = 0.045*). For ASD group, the between GFBs pattern, which was common in both SWP and SWN, was reduction of small-world property by increasing of graph frequencies. For SWP, significant differences were seen between MFB and LFB (*t = −2.72, p = 0.008*), between HFB and MFB (*t = −1.62, p = 0.03*), and between HFB and LFB (*t = −3.4, p = 0.001*). For SWN, significant differences were only seen between MFB and LFB (*t = −2.63, p = 0.01*) and between HFB and LFB (*t = −3.41, p = 0.001*). For TC group, the common pattern was the lower small-world property in the MFB and HFB compared to LFB. For SWP, there were significant difference between MFB and LFB (*t = 1.85, p = 0.05*) and between HFB and MFB (*t = −2.96, p = 0.005*). For SWN, significant were seen between MFB and LFB (*t = −2.76, p = 0.008*) and between HFB and LFB (*t = −3.93, p = 0.0003*).

### 3.2. Results of triadic interactions

For ASD and TC groups, the frequency of triads (|T_i_|), surprise value S, and the ratio of p/p_0_ are listed in Table 2. In all GFBs and for both groups, the unbalanced/balanced triads (T_o_ and T_2_)/(T_1_ and T_3_) were under-presented/over-presented. The under-presented/over-presented shows itself by (p/p_0_ < 1 and S < 0)/(p/p_0_ > 1 and S > 0). The *p*(*T_i_*) is the ratio of the number of triad T_i_ to the total number of triads in the original network.

**Table 2.**
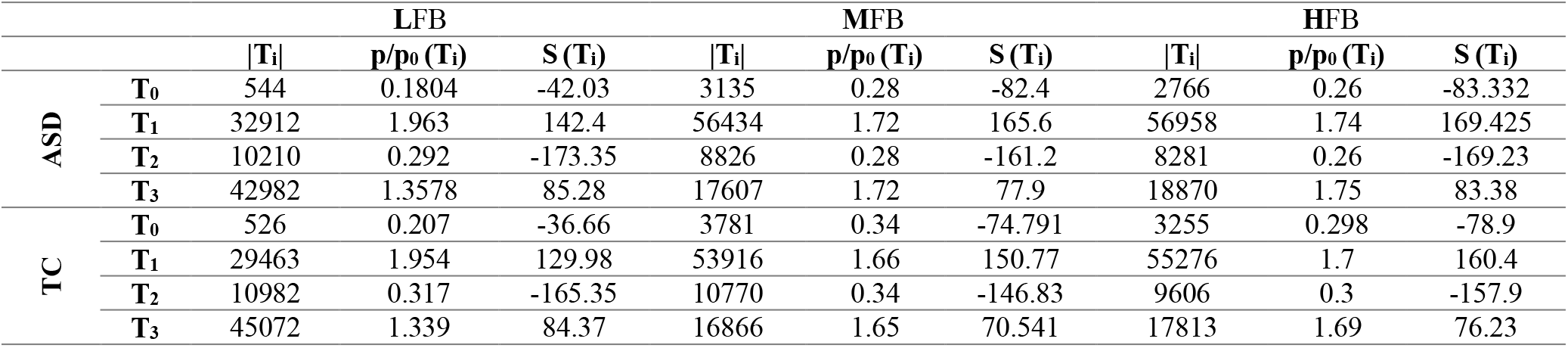
The metrics |T_i_|, S, and the ratio of p/p_0_ averaged over subjects.

The information in this paragraph is for both ASD and TC groups. In all GFBs, the number of balanced triads (|T_1_| and |T_3_|) were more than the number of unbalanced triads (|T_0_| and |T_2_|). In all GFBs, the |T_0_| was the minimum number of triads. In the LFB, the maximum number of triads was devoted to the T_3_. However, in the higher graph frequencies (MFB and HFB), the |T_1_| was the maximum. It should be mentioned that the ratios of |T_1_|/|T_3_| in the MFB and HFB were much bigger than the ratio of |T_3_|/|T_1_| in the LFB.

The results of triadic interaction metrics are shown in Fig 2. For both groups, by moving from LFB to higher graph frequencies, significant high-level changes were seen for |T_0_|, |T_1_|, and |T_3_|. The highest changes were for |T_0_| and |T_3_|. The earlier one increased by a factor of 5-6 and the latter one decreased by a factor of 3. The |T_1_| increased by a factor of close to 2. The significance between-groups differences were found in the MFB (for |T_0_|, |T_1_|, and |T_2_|) and HFB (for |T_3_|). The number of unbalanced triads T_0_ and T_2_ were significantly lower for ASDs compared to TCs in the MFB (*t = −2.03, p = 0.02* for T_0_; *t = −2.12, p = 0.02* for T_2_). In contrast, the |T_1_| and |T_3_| were significantly higher for ASDs compared to TCs in the MFB *(t = 1.89, p = 0.03)* and HFB *(t = 2.02, p = 0.01),* respectively. The general pattern in the MFB and HFB was a higher/lower number of balanced/unbalanced triads for ASDs compared to TCs.

**Fig. 2.**
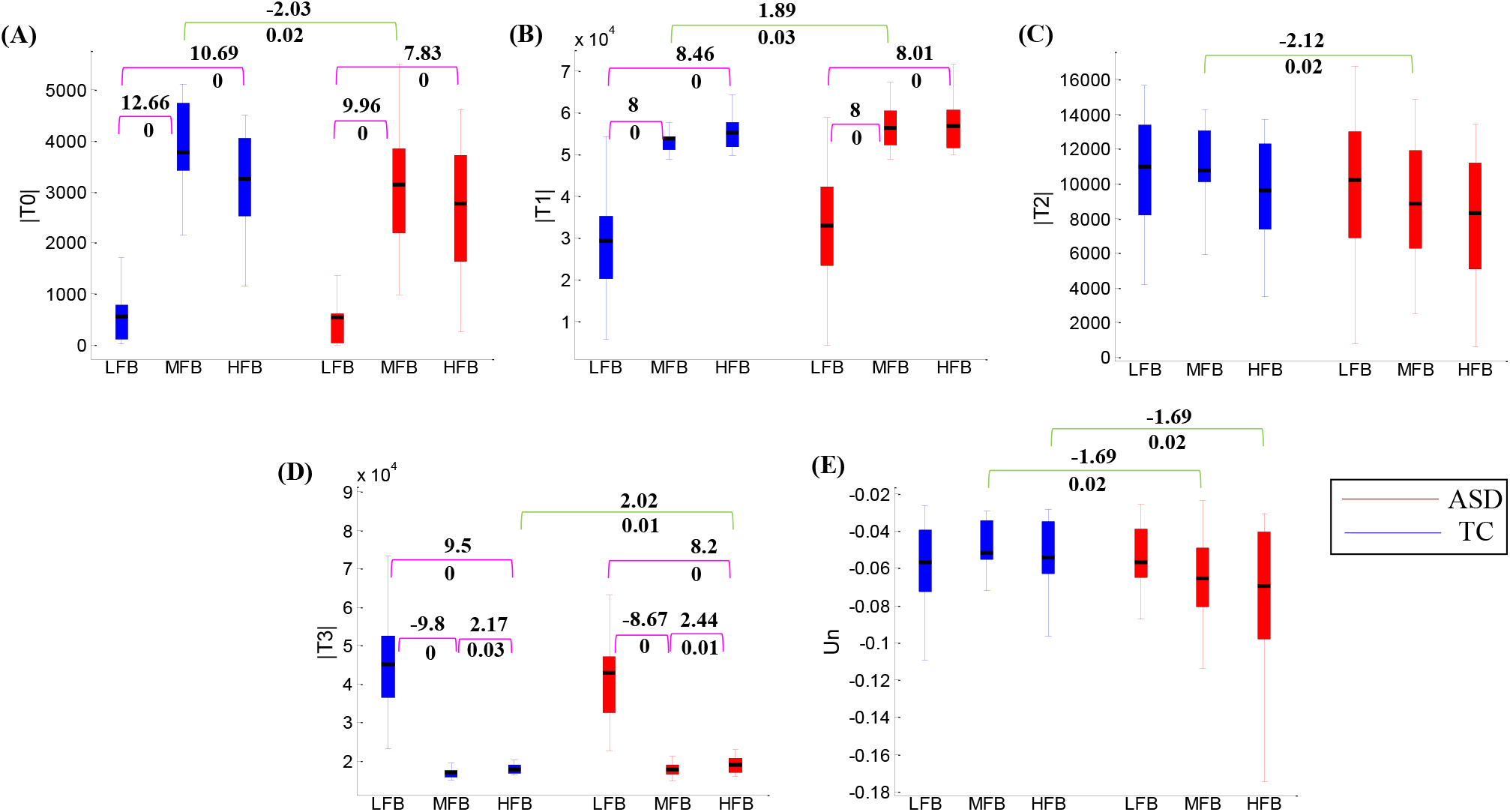
Comparing the triadic interaction metrics between groups and between graph frequency bands. The results are for **(A)** |T_0_|, **(B)** |T_1_|, **(C)** |T_2_|, **(D)** |T_3_|, and **(E)** Un metrics. The green and pink color lines are used to show the results of between groups and between frequency bands, respectively. The values above and under the lines are statistical *t* and corresponding corrected *p* values, respectively. The aforementioned values are only provided for significant results (*p < 0.05*).

The **energies** (Un) of the two groups were negative and approximately equal to each other in the LFB. For TCs, the energies in the MFB and HFB were more than the energy in the LFB. In contrast, increasing the graph frequencies was accompanied by decreasing the energy level for ASDs. Consequently, significant differences were seen between energies of ASD and TC groups in the MFB *(t = −1.69, p = 0.02)* and HFB *(t = −1.69, p = 0.02).*

## 4. Discussion

In this paper, DTI and rfMRI data of SDSU and NYU1 datasets of ABIDE II were used to perform the GSP. Some global metrics of brain functional graph and some metrics of brain functional triadic interactions were studied in the LFB, MFB, and HFB of brain structural graph. Analyzing functional data in structural frequency bands was achieved through the use of GSP. By doing so, some significant differences were found between ASDs and TCs and between GFBs. The main finding was that all significant differences between ASDs and TCs were seen in the MFB and HFB.

The average clustering coefficient of a network is a direct metric for measuring the local specialization of the network (Bullmore and Sporns, 2009). Thus, the results of the clustering coefficient in the MFB may show a decrease of brain functional specialization in ASDs due to an abundance of functional connections (increased clusters of local connections in ASDs). Similar results have also been reported in the DTI study (Li et al., 2018) and the electroencephalography (EEG) study in the theta, beta, and low gamma bands (Ye et al., 2014). The results of clustering coefficient and strength metrics show the overconnectivity in ASDs compared to TCs in the MFB and HFB (Ye et al., 2014; Supekar et al., 2013; Hull et al., 2016; Di Martino et al., 2013). Increased strength and local connectedness in ASDs may suggest impaired network refinement in high-frequency bands of the structural graph (Ye et al., 2014). The results of efficiency show that the global integration of ASDs is more than TCs in the MFB and HFB (Rudie et al., 2013; Li et al., 2018; Ye et al., 2014). Increased clustering coefficient and efficiency indicate increased “cliquishness” properties of functional networks of ASDs in the MFB and HFB, supporting the idea of disrupted balance between global integration and local specialization in ASDs. Similar results have also been seen in theta, beta, low, and high gamma bands in an EEG study (Ye et al., 2014). Both ASD and TC groups had small-world properties (*σ* > 1, *ϕ* > 0.4). However, the ASDs took lower *σ* and significantly lower *φ* than TCs in the HFB. These results show the reduction of optimal balance between network segregation and integration in ASDs.

In this study, Heider’s balance theory was seen for ASD and TC groups in all GFBs (Heider, 1982). This meant that all balanced/unbalanced triads were over-presented/under-presented in the brains of ASDs and TCs in all GFBs (Table 1). However, as has been reported by Moradimanesh and colleagues (Moradimanesh et al., 2021) and some studies of social networks (Leskovek et al., 2010; Szell et al., 2010), T0 may show over-presented behavior in some cases. Thus, this issue was separately investigated on ASDs and TCs in the LFB, MFB, and HFB. Only 3.84% of ASD subjects showed over-presented T_0_ in the LFB. However, this percent value is small and the dominance behavior for T_0_ is under-presentation in the ASDs.

The unbalance triads inject energy into the brain and excite it for changing its state, which in turn results in brain dynamism. (Heider, 1982; Moradimanesh et al., 2021). Thus, the unbalanced triads play a crucial role in the dynamism of the brain, although they under-presented. In this study, the numbers of unbalanced triads T_0_ and T_2_ were significantly lower in ASDs compared to TCs in the MFB. Also, the correlations of |T_0_| and |T_2_| with Autism Diagnostic Interview restricted and repetitive behavior (ADIRRB) score were computed. Negative associations with ADIRRB were seen for both of |*T*_0_| (*r (46) = −0.37; p = 0.01*) and |T_2_| (*r (46) = −0.31; p = 0.036*) (the scores of 46 subjects were available). As a result, it can be said that the lack of enough numbers of unbalanced triads in the MFB leads to difficulty in dynamic switching of the brain which in turn may be a reason for increased repetitive and restricted behaviors in ASDs.

The role of balanced triads from the perspective of structural balance theory is to provide connected modularity. This means that balanced triads provide connected clusters of nodes for the brain in its stable state (Moradimanesh et al., 2021). Thus, the role of balanced triads is important in balancing the functional specialization and integration of brain networks. In this study, the numbers of balanced triads T_1_ and T_3_ were significantly larger for ASDs compared to TCs in the MFB and HFB, respectively. Surely, these differences result in an imbalance of brain networks integration and specialization. These findings are in line with the results of the clustering coefficient and global efficiency in which decreased local specialization and increased global integration were seen in the MFB and HFB.

For ASDs, both having lower numbers of unbalanced triads and having higher numbers of balanced triads can be reasons for having lower energies compared to TCs in the MFB and HFB. This means that lower energies of the ASD group can show both or each of the lower dynamism and unbalanced functional integration and specialization of the brain. To clarify, the correlations of Un with ADIRRB, clustering coefficient, and global efficiency were computed in the MFB and HFB. The correlations with ADIRRB offered (*r (46) = −0.16, p = 0.29*) and (*r (46) = −0.07, p = 0.64*) in the MFB and HFB, respectively. The correlations with clustering coefficient and global efficiency were significant in both MFB and HFB (*all r < −0.9, all p < 1e-18*). As a result, it can be said that lower energies of ASDs are due to having more balanced triads. In another point of view, lower energies of ASDs indicate the disruption of the optimum balance between functional integration and local specialization in the MFB and HFB.

Most of the significant results of this paper were found in the MFB and HFB. Accordingly, the belief that only the first several graph frequency modes contain the most significant information, just as the classical Fourier frequency, (Wang et al., 2018) cannot be valid (Huang et al., 2018), at least for comparison analysis of rfMRI data of ASD subjects and TCs. From another perspective, given the frequency illustrated in Fig 1, it is obvious that the rfMRI data cannot be similar (with the same sign and very low variability) for all connected ROIs. In other words, the brain functional data also exists in the MFB and HFB of the structural graph.

The scenarios of this paper were also implemented using Schaefer atlas with 200 ROIs (FA200) to investigate the reproducibility of results with different numbers of ROIs. For all metrics, results of FA200 were similar to those of FA100 and the significant results were repeated. For interested readers, the results regarding 200 ROIs are represented in S.1 of the supplementary file. By reviewing the results of Tables S1 and 2, it was seen that the signs of statistical values (*t*) were only preserved in the MFB and HFB while changing the number of ROIs. This may be interpreted as the higher reproducibility of results in the MFB and HFB compared to LFB.

Community detection analysis was also performed using the Louvain method in the GraphVar software (Blondel et al., 2008; Kruschwitz et al., 2015). The more detailed results are given in section S.3 of the supplementary file. In each GFB, three modules were found for both ASDs and TCs (Fig. S1). There were found no significant differences between ASDs and TCs when investigating the maximum modularity (*Q*), normalized mutual information (*MIn*), and normalized variation of information (*VIn*) metrics (these metrics are measures of the modular organization of brain networks (more explanations are in S.2 of supplementary file)). However, the *VIn* was much more than *MIn* in the HFB and, particularly, in the MFB. This meant that modules of ASDs and TCs had less common information and more variation to each other in the MFB and HFB. The modularity overlap information confirmed the *VIn* and *MIn* results (Table S5 and 6). These results were consistent with our main finding that the significant differences between ASDs and TCs existed in the MFB and HFB when analyzing the graph global metrics and triadic interaction metrics.

In the community analysis, the ROIs were investigated for translational and local connectivity hub roles. Transitional ROIs facilitate functional integration between modules. Hub ROIs of a module has many connections with other ROIs of that module i.e., they are the core components of that module. In the LFB, the translational roles of the right hemisphere somatomotor 7 and 8 were significantly more for TCs compared to ASDs (transitional ROIs facilitate functional integration between modules). In the MFB and HFB, there was were no ROIs with significant differences between ASDs and TCs in terms of translational role. For hub role, the left hemisphere lateral prefrontal cortex (salience/ventral attention network) in the LFB and the right hemisphere prefrontal cortex 6 (ROI of default mode network) and visual 1 in the MFB were for ASDs. For TCs, the left hemisphere precuneus posterior cingulate cortex 1 (ROI of default mode network) in the MFB and right hemisphere visual 7 were the local hub.

Although the ExploreDTI software was used herein, given its well-established reputation, fiber tractography, in general, has limitations in terms of false positives (and negatives). For insurance, the tractogram of each subject was filtered by linear fascicle evaluation (LiFE) procedure (Pestilli et al., 2014). Then, the SC matrix of filtered tractogram was computed as explained in subsection 2.2.2. For all subjects and atlases, the Pearson correlation analysis between original and filtered SC matrices showed no significance difference (all r > 0.76, all p << 0.0001). The general pattern of scenario results didn’t show any changes, i.e., significant/non-significant results stayed significant/non-significant. Only some changes were seen in statistical *t* values and corresponding *p-values*. Thus, original and corrected SC matrices confirmed each other and were not discrepant each other.

### 4.1. Limitations and future directions

In this paper, only male subjects with a broad age range (5.32-26.6 years) were studied. For the future, the data of both males and females with a narrower age range can be used to provide results for both groups and between genders.

In this study, the underlying graph of GSP was attained using the SC of DTI data and the graph signal was rfMRI data. However, it may be more convenient to employ GSP using only one modality. One can use the FC of rfMRI as the underlying graph of GSP and investigate the reproducibility of the results of this study. Such investigation and finding the proper FC can be a topic for future work. The FC matrices are better to be sparse to reduce sensitivity not only to inter-individual differences in noise but also to global group connectivity differences (e.g., predominant under-connectivity or overconnectivity in ASDs) (Redcay et al., 2013).

In this study, the graph frequencies were divided into three independent bands. For future work, one can divide graph frequencies into more GFBs. This may offer more discriminative features between ASDs and TCs.

The ASD researches dealing with graph global metrics are very limited. One reason might be that the researchers have not found these metrics as significant discriminative features. However, this study demonstrated that using GSP could make these metrics such discriminative features when using them in the MFB and HFB. For future work, graph global and triadic interaction metrics in the MFB and HFB can be used to classify ASDs and TCs. More investigation of these features using machine learning techniques and data of much more subjects may lead to introducing some of the studied metrics as biomarkers.

For both ASD and TC groups, the |T_1_| showed a significant increase in the MFB and HFB compared to LFB whereas the |T_3_| showed a significant decrease in the MFB and HFB compared to LFB. The weakly balanced triads compared to strongly balanced ones are more prone to become unbalanced triads and, as stated in the fourth paragraph of the discussion, the unbalanced triads are responsible for brain dynamism. Therefore, the resources for being more dynamic are more in the MFB and HFB. It might be guessed that having such high resources shows the readiness of the brain for change and adaptation over time (age and neuroplasticity (Voss et al., 2017)). Investigation about this guess can be a topic for future work.

The community structure analysis revealed three modules (sub-networks) for ASDs and TCs in each of the studied GFBs. For future work, the studied metrics can be employed to compare ASDs with TCs at these sub-networks.

## 5. Conclusion

Using GFBs revealed significant differences between ASDs and TCs. The main finding was that the significant differences between ASDs and TCs existed in the MFB and HFB of the structural graph when analyzing the global metrics of the functional graph and triadic interaction of the brain. ASDs in comparison to TCs had significant higher clustering coefficient in the MFB, higher efficiencies in the MFB and HFB, higher strengths in the MFB and HFB, lower SWP in the HFB, lower |T_0_| in the MFB, higher |T_1_| in the MFB, lower |T_2_| in the MFB, higher |T_3_| in the HFB, and lower energies in the MFB and HFB. The results of all scenarios confirmed that using both structural and functional data through GSP tools can open a new reliable avenue to study ASD.

## Supporting information

Supplementary file of paper

## Conflict of Interest

The authors declare no conflict of interest.

## Footnotes

1 http://www.fil.ion.ucl.ac.uk/spm/.

2 https://afni.nimh.nih.gov.

## Notes

### Competing Interest Statement

The authors have declared no competing interest.

