## Supplementary file of paper for "A Comparison Study between Autism Spectrum Disorder and Typically Control in Graph Frequency Bands Using Graph and Triadic Interaction Metrics"

### S.1. Supplementary information of FA200

The results of FA100 are also provided for comparison purposes.

#### **S.1.1.** Results of graph global metrics

Table S1. Results are for FA200. **The metrics assortativity, clustering coefficient, efficiency, radius, diameter, strength, SWP, and SWN averaged over subjects**. The *t* and *p* are statistical and probability values, respectively.

| | **t (p)** | | |
|---|---|---|---|
| | **LFB** | **MFB** | **HFB** |
| **Assortativity** | 0.48 (0.66) | 0.09 (0.93) | -0.25 (0.78) |
| **Clustering coefficient** | -0.56 (0.59) | 2.3 (**0.03**) | 0.74 (0.47) |
| **Efficiency** | -0.42 (0.64) | 2.3 (**0.03**) | 2.1 (**0.04**) |
| **Radius** | 0.38 (0.68) | -0.7 (0.4) | -0.31 (0.73) |
| **Diameter** | -0.21 (0.83) | 1 (0.37) | 0.07 (0.96) |
| **Strength** | 0.06 (0.95) | 2.22 (**0.034**) | 2.15 (**0.038**) |
| **SWP** | -0.44 (0.68) | 0.35 (0.7) | -2.25 (**0.03**) |
| **SWN** | -0.67 (0.5) | -0.9 (0.38) | -1.1 (0.19) |

Table S2. Results are for FA100. **The metrics assortativity, clustering coefficient, efficiency, radius, diameter, strength, SWP, and SWN averaged over subjects**. The *t* and *p* are statistical and probability values, respectively.

| | **t (p)** | | |
|---|---|---|---|
| | **LFB** | **MFB** | **HFB** |
| **Assortativity** | 0.27 (0.82) | 0.48 (0.63) | -0.25 (0.8) |
| **Clustering coefficient** | 0.22 (0.83) | 2.11 (**0.04**) | 1.7 (0.09) |
| **Efficiency** | -0.001 (1) | 2.11 (**0.04**) | 1.93 (**0.04**) |
| **Radius** | -0.11 (0.94) | -0.36 (0.72) | -0.47 (0.69) |
| **Diameter** | 0.73 (0.46) | 0.73 (0.5) | 1.3 (0.21) |
| **Strength** | 0.06 (0.96) | 1.94 (**0.047**) | 1.93 (**0.049**) |
| **SWP** | 0.75 (0.44) | 0.68 (0.46) | -1.98 (**0.045**) |
| **SWN** | -1.38 (0.15) | -1.15 (0.25) | -1.38 (0.165) |

#### **S.1.2.** Results of triadic interaction metrics

Table S3. Results are for FA200. **The metrics $|T_i|$, S, and the ratio of $p/p_0$ averaged over subjects**.

| | | LFB | | | MFB | | | HFB | | |
|---|---|---|---|---|---|---|---|---|---|---|
| | | **$\|T_i\|$** | **$p/p_0$ ($T_i$)** | **S ($T_i$)** | **$\|T_i\|$** | **$p/p_0$ ($T_i$)** | **S ($T_i$)** | **$\|T_i\|$** | **$p/p_0$ ($T_i$)** | **S ($T_i$)** |
| **ASD** | **$T_0$** | 36707 | 0.19 | -98.56 | 24639 | 0.32 | -202.51 | 25221 | 0.33 | -197.07 |
| | **$T_1$** | 207565 | 2.16 | 343.73 | 385183 | 1.68 | 412.34 | 379346 | 1.66 | 402.55 |
| | **$T_2$** | 76818 | 0.31 | -437.17 | 71939 | 0.32 | -407.366 | 75472 | 0.33 | -400.23 |
| | **$T_3$** | 320695 | 1.39 | 223.3 | 124226 | 1.68 | 197.66 | 124885 | 1.66 | 194.4 |
| **TC** | **$T_0$** | 36638 | 0.29 | -85.28 | 24930 | 0.32 | -201.21 | 26046 | 0.34 | -194.92 |
| | **$T_1$** | 186892 | 2.17 | 327.94 | 343499 | 1.67 | 408.84 | 376785 | 1.66 | 397.25 |
| | **$T_2$** | 75333 | 0.32 | -421.49 | 73215 | 0.33 | -404.23 | 77258 | 0.34 | -394.01 |
| | **$T_3$** | 343434 | 1.36 | 224.3 | 124306 | 1.67 | 196.1 | 123573 | 1.65 | 191.22 |

Table S4. Results are for FA100. **The metrics $|T_i|$, S, and the ratio of $p/p_0$ averaged over subjects**.

| | | LFB | | | MFB | | | HFB | | |
|---|---|---|---|---|---|---|---|---|---|---|
| | | **$\|T_i\|$** | **$p/p_0$ ($T_i$)** | **S ($T_i$)** | **$\|T_i\|$** | **$p/p_0$ ($T_i$)** | **S ($T_i$)** | **$\|T_i\|$** | **$p/p_0$ ($T_i$)** | **S ($T_i$)** |
| **ASD** | **$T_0$** | 544 | 0.1804 | -42.03 | 3135 | 0.28 | -82.4 | 2766 | 0.26 | -83.332 |
| | **$T_1$** | 32912 | 1.963 | 142.4 | 56434 | 1.72 | 165.6 | 56958 | 1.74 | 169.425 |
| | **$T_2$** | 10210 | 0.292 | -173.35 | 8826 | 0.28 | -161.2 | 8281 | 0.26 | -169.23 |
| | **$T_3$** | 42982 | 1.3578 | 85.28 | 17607 | 1.72 | 77.9 | 18870 | 1.75 | 83.38 |
| **TC** | **$T_0$** | 526 | 0.207 | -36.66 | 3781 | 0.34 | -74.791 | 3255 | 0.298 | -78.9 |
| | **$T_1$** | 29463 | 1.954 | 129.98 | 53916 | 1.66 | 150.77 | 55276 | 1.7 | 160.4 |
| | **$T_2$** | 10982 | 0.317 | -165.35 | 10770 | 0.34 | -146.83 | 9606 | 0.3 | -157.9 |
| | **$T_3$** | 45072 | 1.339 | 84.37 | 16866 | 1.65 | 70.541 | 17813 | 1.69 | 76.23 |

### S.2. Results of community structure analysis

Community detection analysis was performed using the Louvain method in the GraphVar software. In this analysis, the brain is divided into non-overlapping groups of ROIs so that the numbers of within-group edges are maximum and the numbers of between-group edges are the minimum ones. In this study, the maximum modularity ($Q$), normalized mutual information (*MIn*), normalized variation of information (*VIn*), Classification consistency ($z$), and Classification diversity ($h$) were computed as measures of modular organization. The modularity quantifies how well the graph is divided into subgroups. The *MIn*/*VIn* quantifies how much information is shared/varied by/between the two (different) partitions Ci and Cj of a given network. The $z$ quantifies the degree to which each region is classified in the same module across participants relative to other ROIs in the same module. Brain regions with high $z$ values represent core components of their module and thus act as **local connectivity hubs**. The $h$ quantifies the variability of each region's modular assignment across participants. Regions with high $h$ have a relatively equal probability of being classified into different modules across participants because their connectivity is dispersed between modules from individual to individual. These regions, therefore, represent **transitional ROIs** that facilitate functional integration between modules.

In each GFB, three modules were found for both groups (Fig S1). The $Q$s of ASDs and TCs were 0.61 and 0.65 in the LFB, 0.43 and 0.6 in the MFB, and 0.58 and 0.47 in the HFB, respectively. There were found no significant differences between $Q$s of ASDs and TCs in the LFB ($p = 0.55$), MFB ($p = 0.22$), and HFB ($p = 0.27$), respectively. The (*MIn, VIn*) were (0.49, 0.24) in the LFB, (0.06, 0.43) in the MFB, and (0.16, 0.39) in the HFB, respectively. The (*MIn*, *VIn*) were not significant in the LFB (0.25, 0.25), MFB (0.21, 0.49), and HFB (0.75, 0.31), respectively.

The results of $z$ and $h$ are plotted in Fig S2. In the LFB, the differences of $h$ between ASDs and TCs were significant for **right hemisphere somatomotor 7 and 8** ($p = 0.03$ and $p = 0.01$). The $h$ values of these regions were 0.79 and 0.72 for ASDs and 0.94 and 0.91 for TCs. For $z$ metric, the significant difference was seen in the **left hemisphere lateral prefrontal cortex** which is a region of salience/ventral attention network ($p = 0.04$). The $z$ value of this region was 1.26 for ASDs and -0.11 for TCs.

In the MFB, the **prefrontal cortex 6** (ROI of default mode network), **precuneus posterior cingulate cortex 1** (ROI of default mode network), and **visual 1** showed significant *z* difference between ASDs and TCs (*p = 0, p = 0.02, p = 0.01*). The *z* values of these regions were (1.63, -1.2, 1.83) for ASDs and (-1.7, 1.9, -1.8) for TCs, respectively. These regions are in the left hemisphere.

In the HFB, the **right hemisphere visual 7** showed a significant *z* between ASDs and TCs (*p = 0.005*). The z value of this region was -1.5 for ASDs and 2.03 for TCs.

The modularity overlaps are listed in Table S5 and Table S6. The maximum values of overlap were lower in the MFB and HFB compared to the LFB. This meant that the overlaps of one module of ASD/TC with two or all modules of TC/ASD were high in the MFB and HFB. Module3 of ASD/TC had a high level of overlap with modules1 and 2 (not mosule3) of TC/ASD in the MFB. All of these results were consistent with *VIn* and *MIn* results. These metrics informed that there were less mutual information and higher variation of information in the HFB and, particularly, in the MFB.

All *p-values* were obtained by permutation. To do this, the group difference of the studied measure was calculated and considered as the original value. Then, labels were permuted across groups and the difference between groups was re-calculated. This approach was repeated 400 times to attain a distribution of group differences in the respective modularity metric. By placing the original value in the random distribution of differences, the *p-value* was calculated for the studied measure.

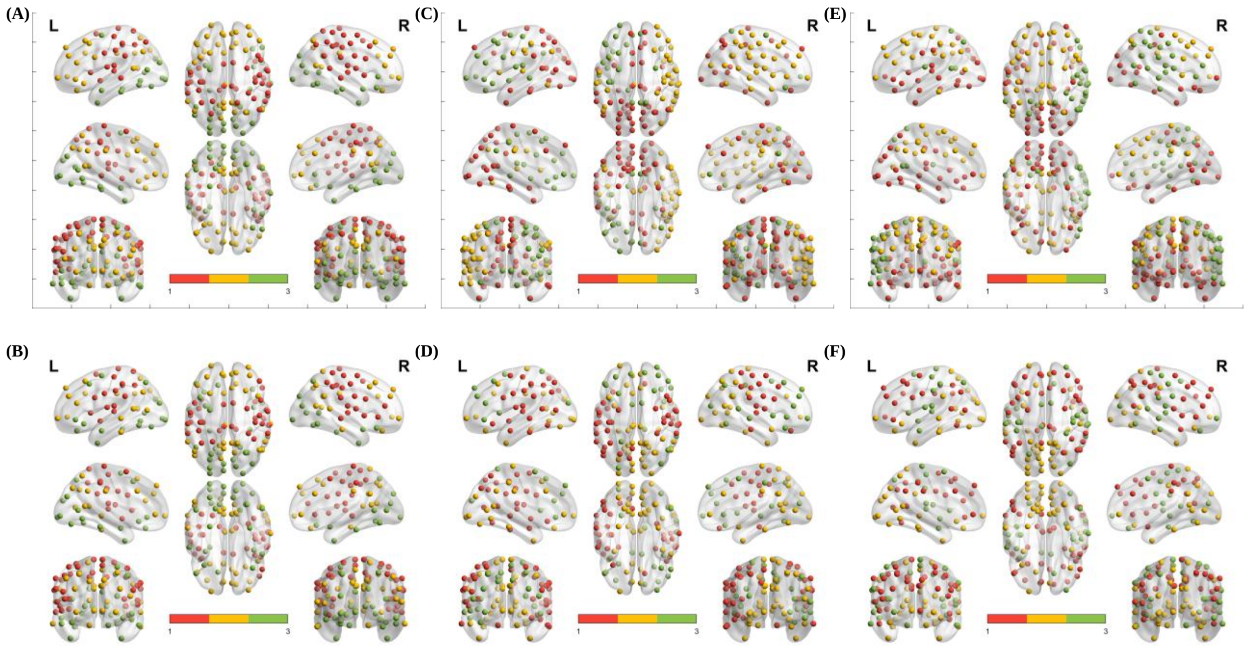


Fig.S1. Modules of ASD and TC groups in the LFB, MFB, and HFB. Results are for **(A)** ASD in the LFB, **(B)** TC in the LFB, **(C)** ASD in the MFB, **(D)** TC in the MFB, **(E)** ASD in the HFB, **(F)** TC in the HFB. The large, medium, and small modules are shown by red, yellow, and green colors, respectively. In the LFB, the colors of the two groups match. In the MFB and HFB, modules of ASDs with red and yellow colors have more overlap with yellow and red color modules of TCs. The visualization is carried out by BrainNet Viewer software.

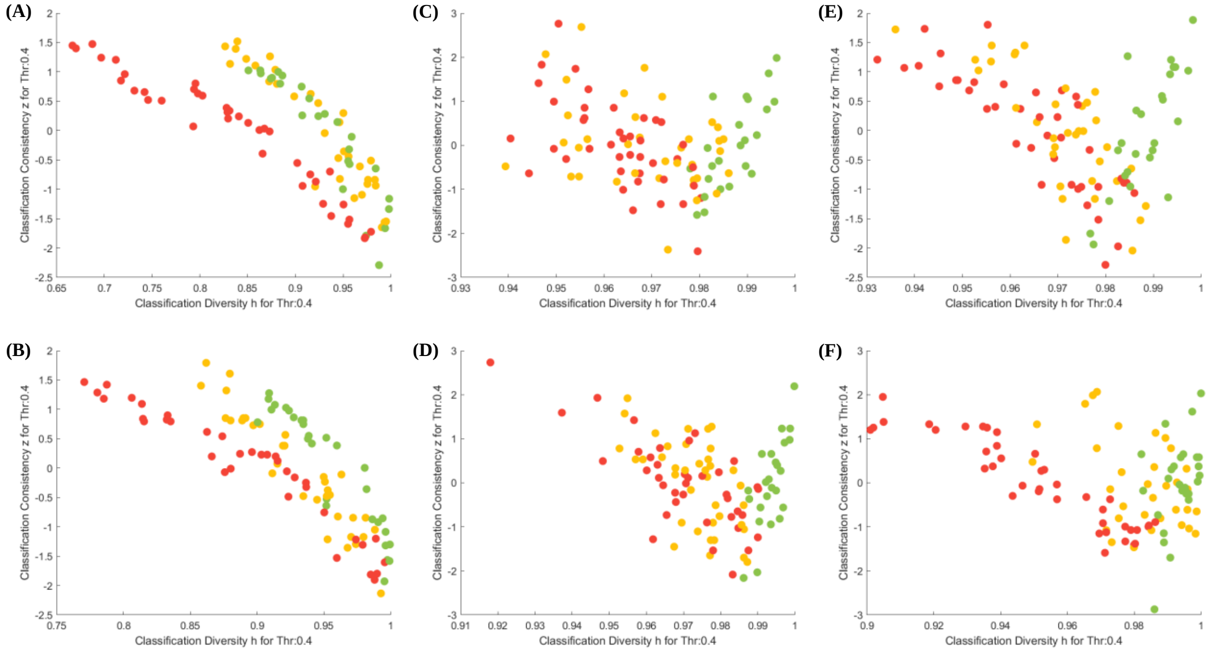


Fig.S2. Results of z versus h are plotted. Results are for **(A)** ASD in the LFB, **(B)** TC in the LFB, **(C)** ASD in the MFB, **(D)** TC in the MFB, **(E)** ASD in the HFB, **(F)** TC in the HFB. The large, medium, and small modules are shown by red, yellow, and green colors, respectively. In the LFB, the colors of the two groups match. In the MFB and HFB, modules of ASDs with red and yellow colors have more overlap with yellow and red color modules of TCs.

Table S5. **The modularity overlap.** Each column represents the overlap of ASD group modules with modules of the TC group. The sum of each column is 1. Mudole1, Module2, and Module3 are the large, medium, and small modules, respectively. In Fig. S1 and 2, these modules are shown by red, yellow, and green colors, respectively.

| | | **ASD** | | | | | | | | |
|---|---|---|---|---|---|---|---|---|---|---|
| | | | **LFB** | | | **MFB** | | | **HFB** | |
| | | Module1 | Module2 | Module3 | Module1 | Module2 | Module3 | Module1 | Module2 | Module3 |
| **TC** | Module1 | **0.825** | 0.0606 | 0.037 | 0.2439 | **0.5278** | **0.3913** | 0.1500 | **0.7143** | 0.3600 |
| | Module2 | 0.15 | **0.7879** | 0.1111 | **0.5610** | 0.1667 | **0.3913** | **0.6000** | 0.1429 | 0.1600 |
| | Module3 | 0.025 | 0.1515 | **0.8519** | 0.2174 | 0.3056 | 0.2174 | 0.2500 | 0.1429 | **0.4800** |

Table S6. **The modularity overlap.** Each column represents the overlap of TC group modules with modules of the ASD group. The sum of each column is 1. Mudole1, Module2, and Module3 are the large, medium, and small modules, respectively. In Fig. S1 and 2, these modules are shown by red, yellow, and green colors, respectively.

| | | **TC** | | | | | | | | |
|---|---|---|---|---|---|---|---|---|---|---|
| | | | **LFB** | | | **MFB** | | | **HFB** | |
| | | Module1 | Module2 | Module3 | Module1 | Module2 | Module3 | Module1 | Module2 | Module3 |
| **ASD** | Module1 | **0.9167** | 0.1714 | 0.0345 | 0.2632 | **0.6053** | 0.3333 | 0.1500 | **0.7273** | 0.3704 |
| | Module2 | 0.0556 | **0.7429** | 0.1724 | **0.5000** | 0.1579 | **0.4583** | **0.6250** | 0.1515 | 0.1852 |
| | Module3 | 0.0278 | 0.0857 | **0.7931** | 0.2368 | 0.2368 | 0.2083 | 0.2250 | 0.1212 | **0.4444** |
